# Satellite glia are essential modulators of sympathetic neuron survival, activity, and autonomic function

**DOI:** 10.1101/2021.09.23.461591

**Authors:** Aurelia A. Mapps, Erica Boehm, Corrine Beier, William T. Keenan, Jennifer Langel, Michael Liu, Samer Hattar, Haiqing Zhao, Emmanouil Tampakakis, Rejji Kuruvilla

**Affiliations:** Department of Biology, Johns Hopkins University, 3400 N. Charles St, 227 Mudd Hall, Baltimore, Maryland 21218, USA; Section on Light and Circadian Rhythms (SLCR), National Institute of Mental Health, NIH, Bethesda, Maryland 20892, USA; Department of Neuroscience, The Scripps Research Institute, La Jolla, California 92037, USA; Department of Medicine, Division of Cardiology, Johns Hopkins University, 720 Rutland Ave, Ross 835, Baltimore, Maryland 21205, USA

## Abstract

Satellite glia are the major glial cells in sympathetic ganglia, enveloping neuronal cell bodies. Despite this intimate association, how satellite glia contribute to sympathetic functions remain unclear. Here, we show that satellite glia are critical for metabolism, survival, and activity of sympathetic neurons and modulate autonomic behaviors in mice. Adult ablation of satellite glia results in impaired mTOR signaling, soma atrophy, reduced noradrenergic enzymes, and loss of sympathetic neurons. However, persisting neurons have elevated activity, and satellite glia-ablated mice show increased pupil dilation and heart rate, indicative of enhanced sympathetic tone. Satellite glia-specific deletion of Kir4.1, an inward-rectifying potassium channel, largely recapitulates the cellular defects observed in glia-ablated mice, suggesting that satellite glia act in part via extracellular K^+^ buffering. These findings highlight neuron-satellite glia as functional units in regulating sympathetic output, with implications for disorders linked to sympathetic hyper-activity such as cardiovascular disease and hypertension.

## Introduction

The sympathetic nervous system prepares the body for “fight or flight” responses and maintains homeostasis during daily activities such as exercise, digestion, or regulation of body temperature. Post-ganglionic neurons, that reside in sympathetic ganglia and project axons to innervate diverse peripheral organs and tissues, mediate key autonomic effects including cardiac output, metabolism, and immune function (Goldstein, 2013). Satellite glia are the major glial cells in sympathetic ganglia (Hanani, 2010; Mapps et al., 2021), and have a unique architecture in completely enveloping neuronal cell bodies (Hanani, 2010). Each neuron and associated glia are thought to form a discrete structural and functional unit (Hanani, 2010). Despite this intimate association, the essential functions of satellite glial cells in the sympathetic nervous system are vastly under-studied.

Satellite glia have been largely characterized by their distinctive location and morphology in sympathetic ganglia (Hanani, 2010). Like sympathetic neurons, satellite glial cells are derived from multipotent neural crest precursors, and form thin cytoplasmic sheaths around cell bodies, dendrites, and synapses of sympathetic neurons, with only 20 nm of space, the width of a synaptic cleft, between neuronal and glial membranes (Hanani, 2010; Hanani and Spray, 2020; Pannese, 1981). Multiple satellite glia surround a single neuron and are connected with each other, and with neurons, via gap junctions, with the number of glial cells per neuron being positively correlated to soma size (Hanani, 2010; Hanani and Spray, 2020; Ledda et al., 2004). This unique arrangement places satellite glial cells in an ideal position to be critical regulators of neuronal connectivity, synaptic transmission, and homeostasis. Studies in sympathetic neuron-glia co-cultures have suggested roles for satellite glia in promoting dendrite growth, synapse formation, modulating extracellular ion and neurotransmitter concentrations, and regulating synaptic transmission (Enes et al., 2020; Feldman-Goriachnik et al., 2018; Hanani, 2010; Tropea et al., 1988). Satellite glia also envelop neuronal cell bodies in sensory ganglia in the peripheral nervous system (PNS) (Hanani and Spray, 2020). Recent studies implicate sensory satellite glia in regulating chronic pain through modulating neuronal hyper-excitability (Kim et al., 2016), and in promoting axon regeneration after peripheral nerve injury *in vivo* (Avraham et al., 2020). Satellite glia have been proposed to be closest to astrocytes in the central nervous system (CNS) with respect to expression of machinery related to neurotransmitter uptake/turnover, inward-rectifying potassium channels, functional coupling via gap junctions, and close association with synapses (Hanani and Spray, 2020; Hanani and Verkhratsky, 2021). In contrast to the wealth of information on CNS astrocytes and emerging evidence of the significance of sensory satellite glia in the PNS, little is known about the functions of satellite glia in the sympathetic nervous system *in vivo*.

Here, using genetic ablation in mice, we reveal that loss of satellite glia results in impaired metabolic signaling, soma atrophy, reduced expression of noradrenergic enzymes, and enhanced apoptosis of adult sympathetic neurons. The persisting neurons, however, show elevated activity as revealed by increased neuronal c-Fos expression. Consistently, satellite glia-ablated mice had enhanced circulating norepinephrine and elevated heart rate, indicative of heightened sympathetic tone. We further deleted Kir4.1, an inward-rectifying potassium channel, specifically in satellite glia in mice. Satellite glia-specific deletion of Kir4.1, largely recapitulates the cellular phenotypes observed in glia-ablated mice, suggesting that satellite glia support neurons, in part, via extracellular K^+^ buffering. These findings reveal that satellite glia provide critical metabolic and trophic support to sympathetic neurons, and are essential modulators of the ganglionic milieu, neuronal activity, and resulting autonomic behaviors.

## Results

### Inducible ablation of satellite glia using *BLBP:iDTA* mice

Brain Lipid Binding Protein (BLBP), which encodes for a fatty acid transporter, is one of the most highly expressed transcripts in mouse satellite glial cells (Avraham et al., 2020; Kurtz et al., 1994; Mapps et al., 2021). Using immunostaining for BLBP and Tyrosine Hydroxylase (TH), a marker for noradrenergic sympathetic neurons, we observed BLBP-positive satellite glia in the mouse sympathetic ganglia (the Superior Cervical Ganglia or SCG) at both developmental and adult stages (**Figure 1A**). BLBP labeling was “patchy” in the embryonic and neonatal ganglia, a period when satellite glia undergo migration into the ganglia and are also proliferating (Hall and Landis, 1992). However, by two weeks after birth, BLBP-positive satellite glial cells had expanded processes around neuronal cell bodies, and formed thin ring-like glial sheaths around neuronal somas when observed at P31 (**Figure 1A**).

**Figure 1.**
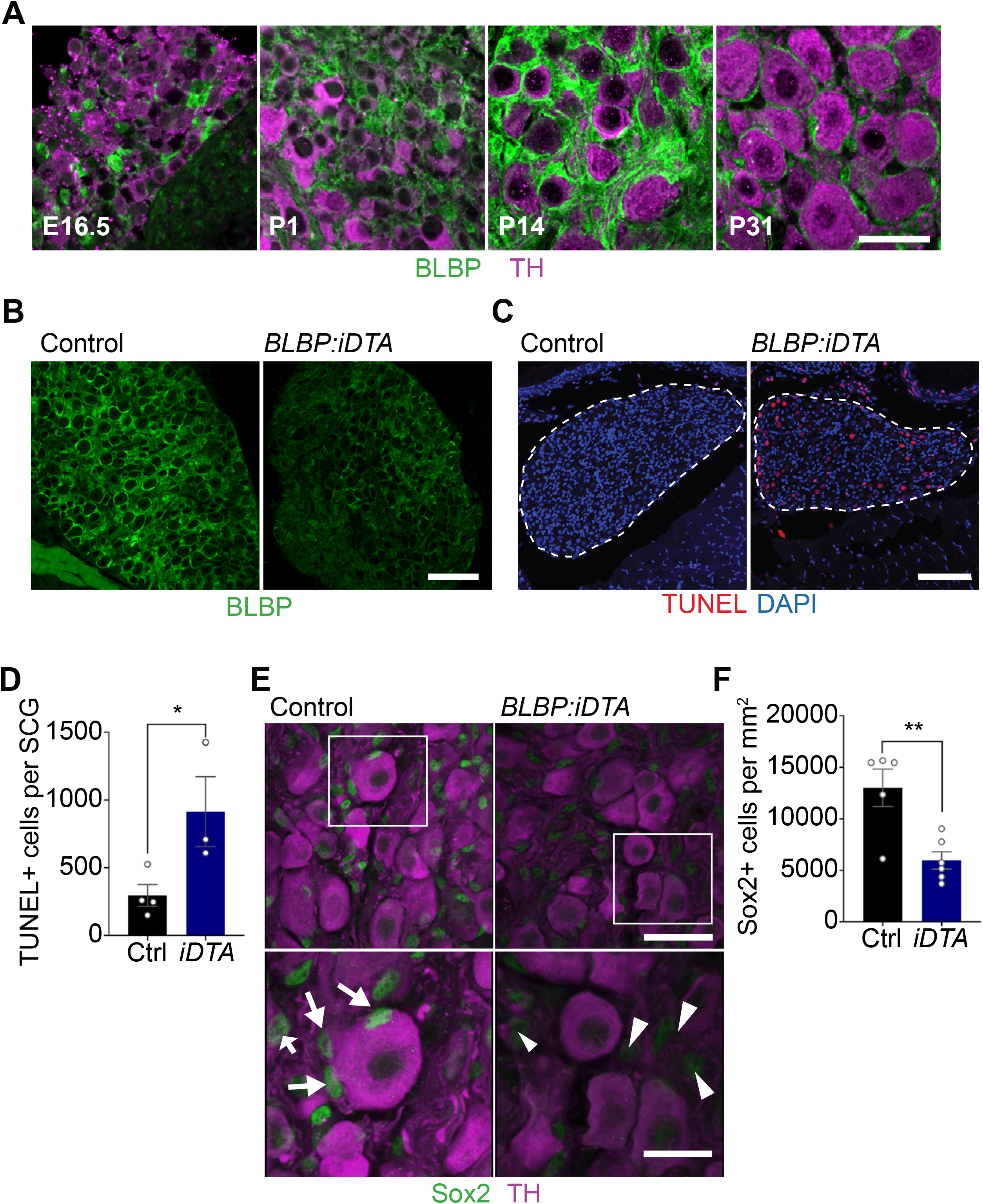
DTA-mediated ablation of satellite glia in sympathetic ganglia. (**A**) Satellite glial cells, immunolabeled with brain lipid binding protein (BLBP, green) progressively ensheathe sympathetic neuron cell bodies, labeled with tyrosine hydroxylase (TH, magenta) in the superior cervical ganglia (SCG) during development. Time-points shown are embryonic day 16.5 (E16.5) and postnatal days P1, P14, and P31. Scale bar: 30μm. (**B**) Reduced BLBP expression in *BLBP:iDTA* SCG relative to control ganglia at 14 days after the last tamoxifen injection. Scale bar:100μm. (**C**) Increased apoptosis in *BLBP:iDTA* SCG (outlined in dashed line) compared to controls as detected by TUNEL labeling (red). Nuclei in tissue sections are labeled with DAPI (blue). Scale bar:100μm. (**D**) Quantification of apoptotic cells in SCGs from n=4 control and 3 mutant mice. Data presented as mean ± s.e.m. *p<0.05, t-test. (**E-F**) Decrease in Sox2-positive (green) satellite glial cells in *BLBP:iDTA* SCG. Arrows indicate Sox2-labeled nuclei of satellite glia associated with sympathetic neuron cell bodies. Arrowheads indicate Sox2-labeled satellite glia that are not associated with neurons. Scale bar:100μm for upper panels, and 30 μm in insets. Data are presented as mean ± s.e.m from n=5 animals per genotype, **p<0.01, t-test.

In the adult PNS, BLBP expression is restricted to satellite glia, and is not detected in Schwann cells based on single-cell RNA sequencing analysis and characterization of *BLBPcre-ER-*driven reporter mice (Avraham et al., 2020). To accomplish ablation of satellite glia, we crossed *BLBPcre-ER* mice (Maruoka et al., 2011) with *ROSA26-eGFP-DTA* mice (Ivanova et al., 2005), where Cre drives expression of a copy of *diphtheria toxin subunit A* (*DTA*). At postnatal day 30, *BLBPcre-ER*;*ROSA26-eGFP-DTA* mice were injected with either vehicle (corn oil) or tamoxifen (180 mg/kg body weight) for 5 consecutive days, and all analyses were performed at 5 or 14 days post injection. By 2 weeks, we observed a drastic loss of BLBP expression in sympathetic ganglia from tamoxifen-treated mice (henceforth referred to as *BLBP:iDTA* mice) compared to vehicle-injected control (Ctrl) mice (**Figure 1B**). While satellite glia in control ganglia formed characteristic ring-like structures around neuronal soma, glial organization was disrupted in mutant mice with diminished BLBP staining detected within the ganglia. To determine if DTA expression results in cell death, we assessed apoptosis using TUNEL labeling and observed a 3-fold increase in TUNEL-positive cells in *BLBP:iDTA* sympathetic ganglia compared to controls (**Figures 1C-D**).

To characterize the loss of satellite glial cells, we performed immunostaining with an antibody against Sox2, a transcription factor expressed in satellite glia (Koike et al., 2014). We observed fewer Sox2-positive glial cells in contact with individual sympathetic neuron cell bodies at 5 days after the last tamoxifen injection (**Figures S1A-B**), although satellite glial numbers appeared to be unaffected at this time point (**Figure S1C**). However, by 14 days post-tamoxifen injection, there was a pronounced loss of glial cells (54.2% decrease) in sympathetic ganglia in *BLBP:iDTA* mice (**Figures 1E-F**). We did not observe increased infiltration by macrophages as assessed by IBA-1 immunohistochemistry (**Figures S1D-E)**, suggesting that satellite glia ablation does not trigger increased inflammation. Further, EdU labeling indicated that satellite glial cells do not undergo increased proliferation as a compensatory or injury-induced response after DTA expression (**Figures S1F-G)**. Together, these results provide evidence in support of DTA-induced depletion of satellite glia, in the absence of inflammation or reactive proliferative changes, in *BLBP:iDTA* sympathetic ganglia.

### Satellite glia depletion impairs noradrenergic enzyme expression, metabolism, and survival of sympathetic neurons

When performing TH immunostaining to visualize sympathetic neuron morphology, we noticed a striking down-regulation of TH immunoreactivity in sympathetic neuronal cell bodies from *BLBP:iDTA* mice. TH is the rate-limiting enzyme in the biosynthesis of Norepinephrine (NE), the classical sympathetic neurotransmitter. Quantitative PCR (qPCR) analyses revealed a drastic down-regulation in *TH* and *Dopamine Beta-Hydroxylase* (DBH) transcript levels, (88% and 99% decrease, respectively) (**Figures 2A-B**). DBH converts dopamine to norepinephrine in the NE biosynthetic pathway. Thus, satellite glia are essential for maintenance of noradrenergic enzymes in sympathetic neurons.

**Figure 2.**
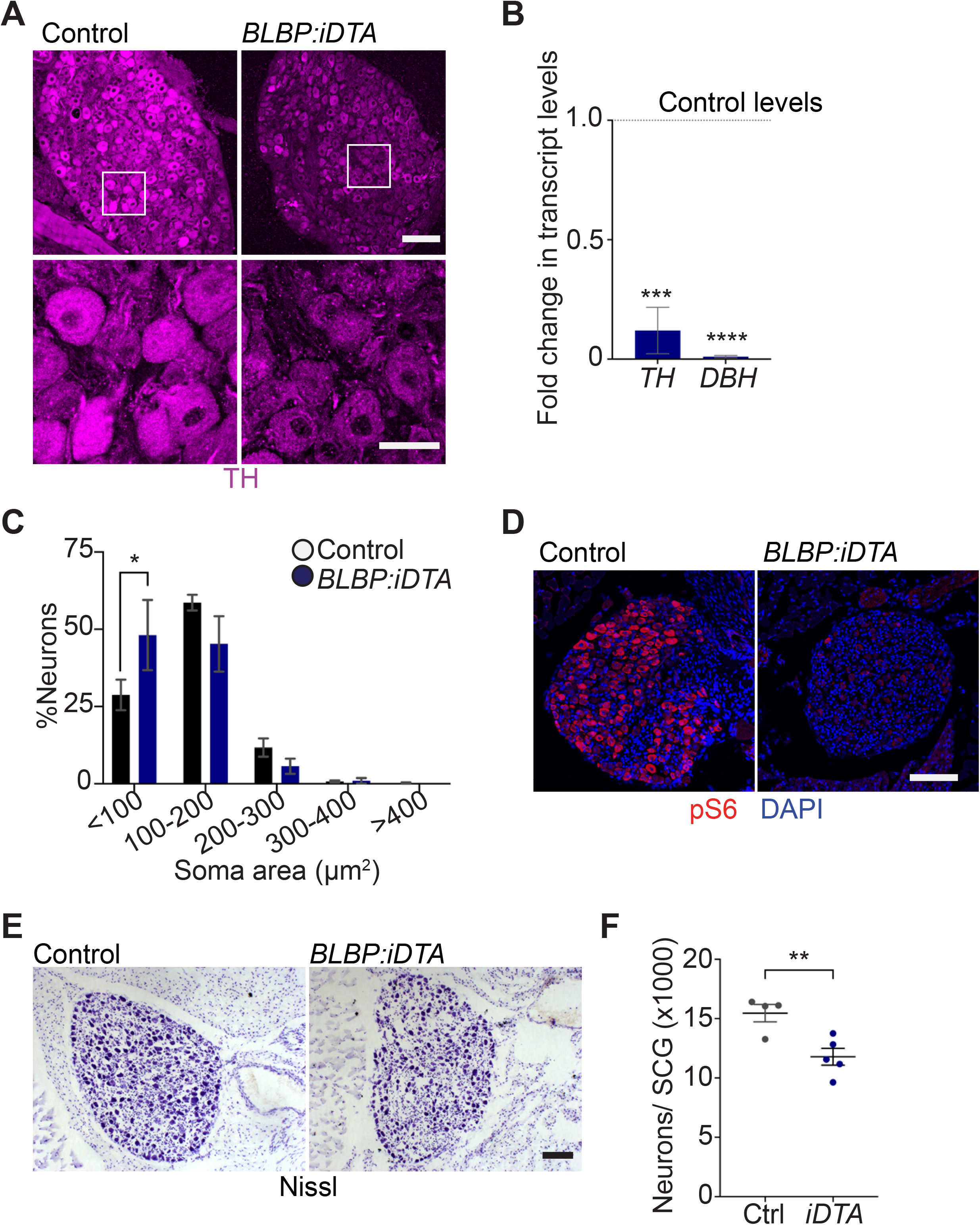
Neuronal defects in NE biosynthesis, metabolism, and survival in satellite glia-depleted mice. **(A)** TH expression is down-regulated in *BLBP:iDTA* sympathetic neurons. Insets also show atrophied neuronal cell bodies in mutant ganglia compared to controls. Scale bar:100μm for upper panels and 30 μm for insets. (**B**) Transcripts for *TH* and *DBH*, key enzymes in norepinephrine biosynthesis, are decreased in *BLBP:iDTA* SCG relative to control ganglia. Data are presented as mean ± s.e.m from SCGs collected from n=3-4 animals per genotype. ***p<0.001, ****p<0.0001, t-test with Bonferroni-Dunn’s correction. (**C**) Histogram shows a greater distribution of smaller soma sizes in mutant neurons compared to controls. Results are mean ± s.e.m from n=5 of animals per genotype, *p<0.05, two-way ANOVA with Bonferroni’s correction. (**D**) pS6 immunostaining indicates diminished mTOR activity in *BLBP:iDTA* SCG relative to control ganglia. Scale bar:100μm. (**E-F**) Cell counts in Nissl-stained SCG tissue sections show reduced sympathetic neuron numbers in satellite glia-depleted mice 14 days after the last tamoxifen injection. Results are mean ± s.e.m from n=4 control and 5 mutant animals. **p<0.01, t-test.

Using TH immunohistochemistry, we also observed pronounced atrophy of neuronal cell bodies in sympathetic ganglia after satellite glia loss (**Figure 2A**). Quantification of soma sizes revealed significantly reduced neuronal soma areas in satellite glia-depleted mice, with a greater distribution of smaller-sized cell bodies in mutant ganglia relative to controls (**Figure 2C**). The average soma area of mutant neurons was 127.5 ± 8 μm^2^, compared to 151.2 ± 16 μm^2^ of control neurons. Since the PI3K/Akt/mTOR pathway is a known regulator of neuronal soma size (Kwon et al., 2006; van Diepen et al., 2009; Zhou et al., 2009), we assessed expression of phosphorylated ribosomal protein 6 (pS6), a key downstream effector of the pathway, using immunostaining. We observed a dramatic reduction in pS6 immunoreactivity in satellite glia-depleted ganglia (**Figure 2D**), suggesting that impaired mTOR signaling might contribute to neuronal atrophy. Given the soma atrophy and that TUNEL-labeled cells also included several cells with large ovoid-shaped nuclei that appeared to be neuronal (**Figure 1C**), we asked if satellite glia ablation affects neuron survival. Indeed, we found that satellite glia depletion results in the loss of 24% of adult sympathetic neurons (15460 ± 734 WT *vs*. 11790 ± 706 mutant neurons) (**Figures 2E-F**), using Nissl staining and cell counts in SCG tissue sections. Despite the decrease in neuronal numbers, sympathetic axon innervation was maintained in target organs in *BLBP:iDTA* mice, when assessed by whole-mount TH immunostaining in iDISCO-cleared tissues and light sheet microscopy (**Figures S2A-C**). Intriguingly, TH levels in mutant axons appeared to be similar to that in controls, despite the marked reduction in *TH* mRNA and protein in neuronal cell bodies residing in the ganglia (**Figure S2A**), suggesting differential regulation of TH distribution in neuronal soma *vs*. axons.

Together, these results indicate that satellite glia are essential for providing trophic and metabolic support to adult sympathetic neurons, and for regulation of the noradrenergic biosynthetic machinery, specifically in neuronal cell bodies.

### Sympathetic activity is elevated in satellite glia-depleted mice

Given neuronal deficits with satellite glia depletion, we sought to determine if autonomic responses were impacted in *BLBP:iDTA* mice. In mammals, pupil size can serve as a non-invasive and rapid readout for autonomic function (McDougal and Gamlin, 2015). Pupil size is modulated by a balance of sympathetic versus parasympathetic activity, with the sympathetic component regulating pupil dilation while parasympathetic activity controls pupil constriction (McDougal and Gamlin, 2015). To measure basal pupil size, control and *BLBP:iDTA* mice were dark-adapted for 2 days, and pupil sizes recorded for 5-10 seconds in the dark in non-anesthetized mice (Keenan et al., 2016). Surprisingly, despite the neuronal loss and down-regulated noradrenergic biosynthetic enzymes in neuronal soma, we observed increased basal pupil areas in *BLBP:iDTA* mice compared to controls (**Figures 3A-B**). To ask whether this phenotype is due to decreased parasympathetic activity, we measured pupil constriction in response to increasing light intensities, ranging from 0.01-1000 lux, administered for 30 seconds. Light onset at 0.1 lux or higher resulted in rapid constriction with greater constrictions at higher light intensities in both *BLBP:iDTA* and control mice (**Figure S3A**). The intensity response curves were virtually identical for the two groups (**Figure S3A**). These results suggest that parasympathetic function is intact in *BLBP:iDTA* mice, and that the enlarged pupil areas likely reflect an increase in sympathetic tone with the loss of satellite glia.

**Figure 3.**
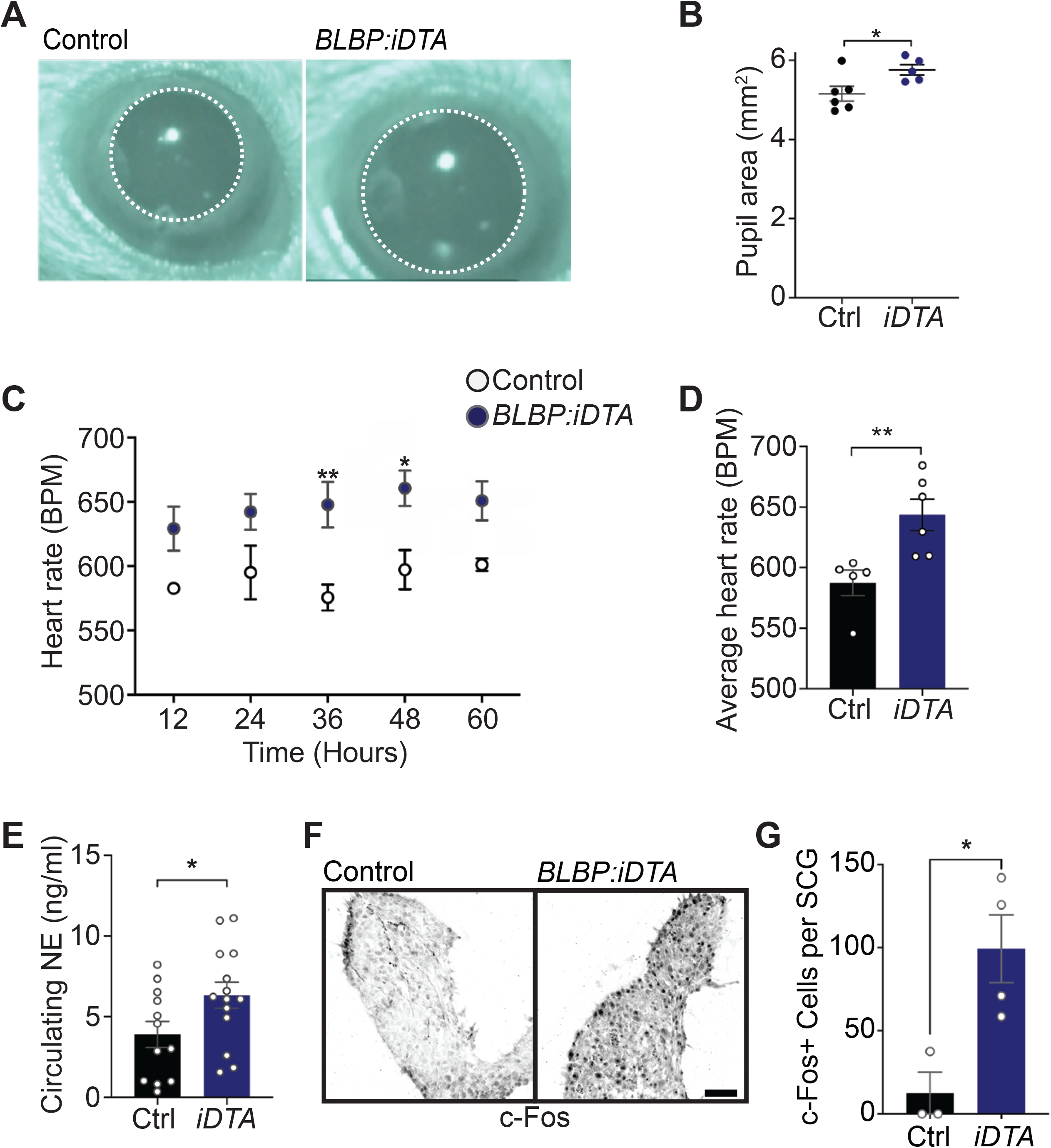
Elevated sympathetic activity in satellite glia-depleted mice. **(A-B)** Dark-adapted *BLBP:iDTA* mice have increased basal pupil size, compared to control litter-mates. Results are presented as mean ± s.e.m from n=6 control and 5 mutant animals. *p<0.05, t-test. (**C**) *BLBP:iDTA* mice exhibit elevated heart rate, relative to controls. ECGs were recorded continuously in conscious mice for 7 days, although only data for 4^th^-7^th^ days after insertion of lead implants are included in the analysis. Results are presented as mean ± s.e.m from n=5 control and 6 mutant animals. *p<0.05, **p<0.01, two-way ANOVA with Bonferroni’s correction. (**D**) Average heart rate over 4^th^-7^th^ days after lead implantation. Results are mean ± s.e.m from n=5 control and 6 mutant animals. **p<0.01, t-test. (**E**) Circulating norepinephrine levels are elevated in *BLBP:iDTA* mice. Results are mean ± s.e.m from n=11 control and 13 mutant animals. *p<0.05, t-test. (**F-G**) Increased c-Fos-positive sympathetic neurons in mutant ganglia compared to controls. Results are mean ± s.e.m from n=3 control and 4 mutant animals. *p<0.05, t-test.

As a second and independent assessment of autonomic function, we measured heart rate and heart rate variability (HRV) using electrocardiogram (ECG) recordings in mice (Thireau et al., 2008). Increased sympathetic activity results in an accelerated heart rate and decreased HRV, defined as the variation in time intervals between consecutive heartbeats (Thireau et al., 2008). Strikingly, *BLBP:iDTA* mice exhibited increased heart rates (**Figures 3C-D**) and decreased HRV (**Figure S3B**) compared to controls, consistent with increased sympathetic tone after two weeks of satellite glia depletion.

To understand the molecular and/or cellular basis for enhanced sympathetic activity in satellite glia-depleted mice, we measured circulating Norepinephrine (NE) levels, and observed a significant 1.6 fold increase in *BLBP:iDTA* mice compared to controls (**Figure 3E**). These results suggest that, despite the down-regulation of NE biosynthetic machinery in neuronal cell bodies (**see Figures 2A-B**), neurotransmitter secretion is augmented in sympathetic axons, and/or its re-uptake/degradation decreased, in satellite glia-depleted mice. NE acts through α- and β-adrenergic receptors in target tissues. Heightened sympathetic activity and increased circulating NE elicits the compensatory down-regulation of adrenergic receptor levels and activities in target tissues (Eschenhagen, 2008). We found significantly decreased expression of *Adrb1*, the major adrenergic receptor for NE signaling in the heart (de Lucia et al., 2018), as well as *Adrb2* and *Adra2c*, using q-PCR analyses of cardiac tissue (**Figure S3C**). Lastly, immunostaining for c-Fos, an immediate early transcription factor that serves as a reporter of neuronal activity (Sheng and Greenberg, 1990), revealed a striking 8-fold increase in the number of c-Fos-positive sympathetic neurons in *BLBP:iDTA* ganglia compared to controls (**Figures 3F-G**).

Together, these results indicate that satellite glia depletion results in elevated sympathetic neuron activity, impaired NE homeostasis, and autonomic behavioral defects in mice.

### Satellite glia-specific deletion of *Kir4*.*1* disrupts sympathetic neuron activity

A key determinant of neuronal excitability is glia-mediated spatial buffering of extracellular potassium ions (K^+^) (Haydon, 2001; Kuffler, 1967). Kir4.1 is a glial-specific, inwardly rectifying K^+^ channel that shows the highest expression in astrocytes and satellite glia (Hanani and Spray, 2020; Olsen et al., 2015; Vit et al., 2008). In mice, global or tissue-specific *Kir4*.*1* deletion results in impaired K^+^ and glutamate homeostasis, loss of glial K^+^ conductance, neuronal excitability, epileptic seizures, and pain-like behaviors (Cui et al., 2018; Djukic et al., 2007; Olsen et al., 2015; Tang et al., 2010; Vit et al., 2008). We observed a pronounced down-regulation of *Kir4*.*1* mRNA in *BLBP:iDTA* sympathetic ganglia, compared to controls, using RNAscope single-molecule Fluorescence *in situ* Hybridization (smFISH) (**Figure S4A**). To ask if impaired K^+^ buffering might contribute to the neuronal defects observed in satellite glia-ablated mice, we generated satellite glia-specific *Kir4*.*1* knockout mice (*Kir4*.*1 cKO*) by crossing *BLBPcre-ER* mice to *Kir4*.*1* floxed mice (Djukic et al., 2007). *BLBPcre-ER;Kir4*.*1*^*fl/fl*^ (henceforth called *Kir4*.*1 cKO*) mice were treated with vehicle or tamoxifen for five consecutive days to conditionally delete Kir4.1 from satellite glial cells. Expression of Kir4.1, visualized by immunofluorescence, was significantly reduced in *Kir4*.*1 cKO* sympathetic ganglia (**Figure 4A**), and qPCR analysis indicated a 76% decrease in *Kir4*.*1* transcript levels (**Figure 4B**).

**Figure 4.**
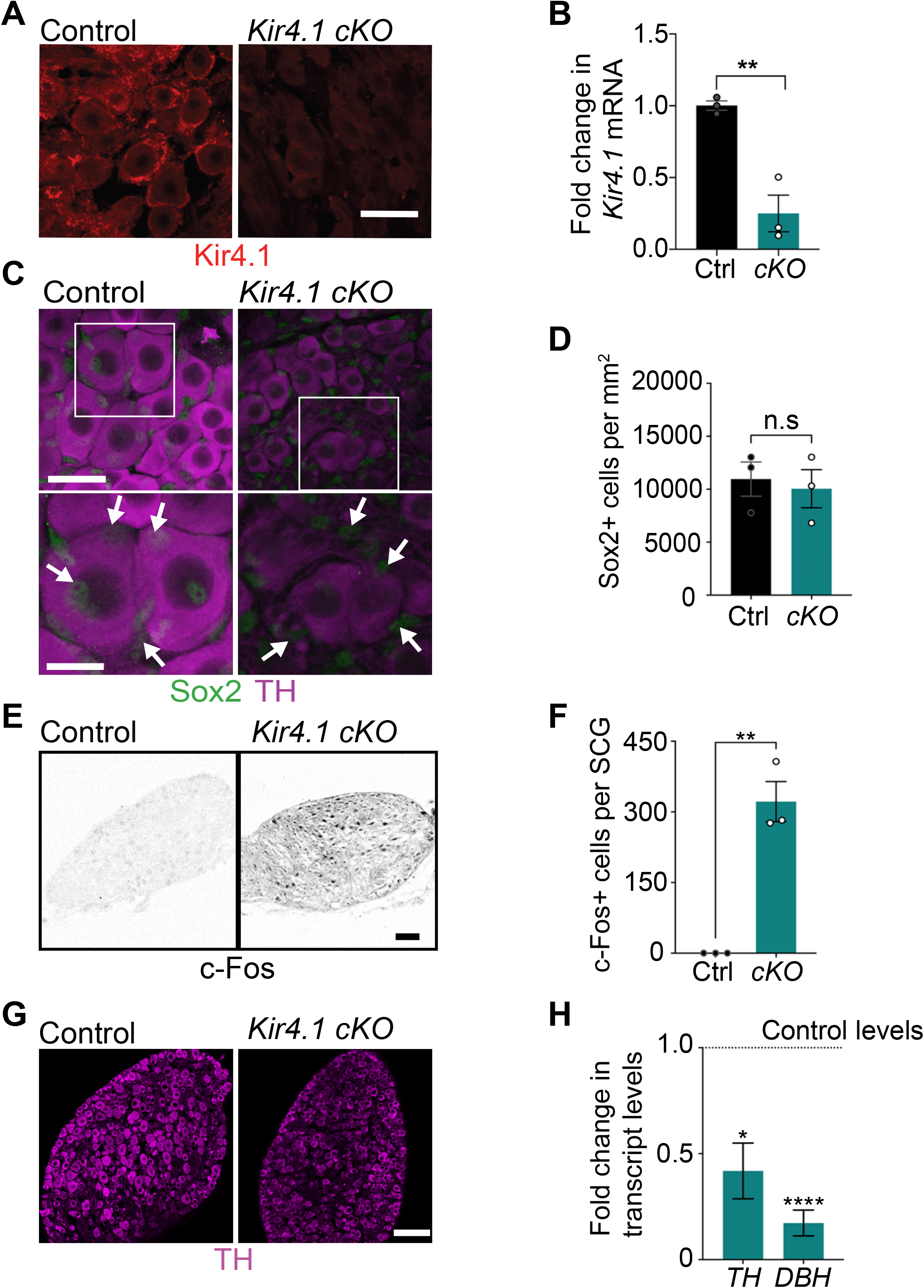
Satellite glia-specific *Kir4*.*1* loss impairs NE enzyme expression and neuron activity. (**A-B**) Reduced Kir4.1 protein and transcript in *Kir4*.*1 cKO* mice. Scale bar: 30μm. Data in (**B**) are mean ± s.e.m from n=3 animals per genotype, **p<0.01, t-test. (**C-D**) Sox2-positive satellite glial cell numbers are unaffected in *Kir4*.*1 cKO* SCG. Arrows indicate Sox2-labeled nuclei of satellite glia associated with sympathetic neuron cell bodies. Arrowheads indicate Sox2-labeled satellite glia that are not associated with neurons. Scale bar:100μm for upper panels and 30 μm in insets. Data are presented as mean ± s.e.m from n=3 animals per genotype, n.s. not significant, t-test. (**E-F**) *Kir4*.*1* deletion in satellite glia results in an increase in c-Fos-positive sympathetic neurons. Scale bar:100μm. Quantifications are mean ± s.e.m from n=3 animals per genotype, **p<0.01, t-test. (**G-H**) Down-regulation of NE biosynthetic enzymes, TH and DBH, in *Kir4*.*1 cKO* sympathetic ganglia. Results are mean ± s.e.m from n=5 animals per genotype, *p<0.05, ****p<0.0001, t-test with Bonferroni’s correction.

Loss of Kir4.1 did not alter satellite glial cell numbers or their overall morphology in sympathetic ganglia as assessed by quantification of Sox2-positive cells and BLBP immunostaining, respectively **(Figures 4C-D, and S4B)**. However, we observed a marked increase in the number of c-Fos-positive cells in *Kir4*.*1 cKO* SCGs compared to controls (**Figures 4E-D**). The majority of c-Fos positive cells in *Kir4*.*1 cKO* ganglia appeared to be neurons based on their morphology. Intriguingly, despite increased c-Fos-positive sympathetic neurons, Kir4.1 deletion resulted in down-regulated expression of noradrenergic biosynthetic enzymes, TH and DBH, in neuronal cell bodies (**Figures 4G-H**), similar to the phenotype observed with satellite glia depletion.

Together, these results indicate that satellite glial cells control sympathetic neuron activity and noradrenergic enzyme expression via Kir4.1-dependent regulation of the neuronal micro-environment.

### Satellite glia-specific deletion of *Kir4*.*1* elicits neuron atrophy and apoptosis

We next addressed the relevance of satellite glia Kir4.1 expression in sympathetic neuron viability. Quantification of soma sizes revealed that neurons undergo atrophy in *Kir4*.*1 cKO* sympathetic ganglia (**Figure 5A**), similar to the defect observed with DTA-induced loss of satellite glia. Consistently, pS6 immunostaining of SCG tissue sections indicated a pronounced decrease in mTOR signaling in mutant ganglia, compared to controls (**Figure 5B**). Notably, we also observed an 8-fold increase in apoptotic cells in *Kir4*.*1 cKO* sympathetic ganglia using TUNEL labeling (**Figures 5C-D**). Since the numbers of Sox2-positive satellite glial cells were unaffected by Kir4.1 deletion (**see Figures 4C-D**), these results imply that the dying cells are primarily sympathetic neurons. Indeed, we observed a 22% decrease in neuronal numbers in *Kir4*.*1 cKO* sympathetic ganglia following Nissl staining and cell counts at two weeks after the last tamoxifen injection (**Figures 5E-F**). These results highlight that Kir4.1 is necessary for the survival of adult sympathetic neurons.

**Figure 5.**
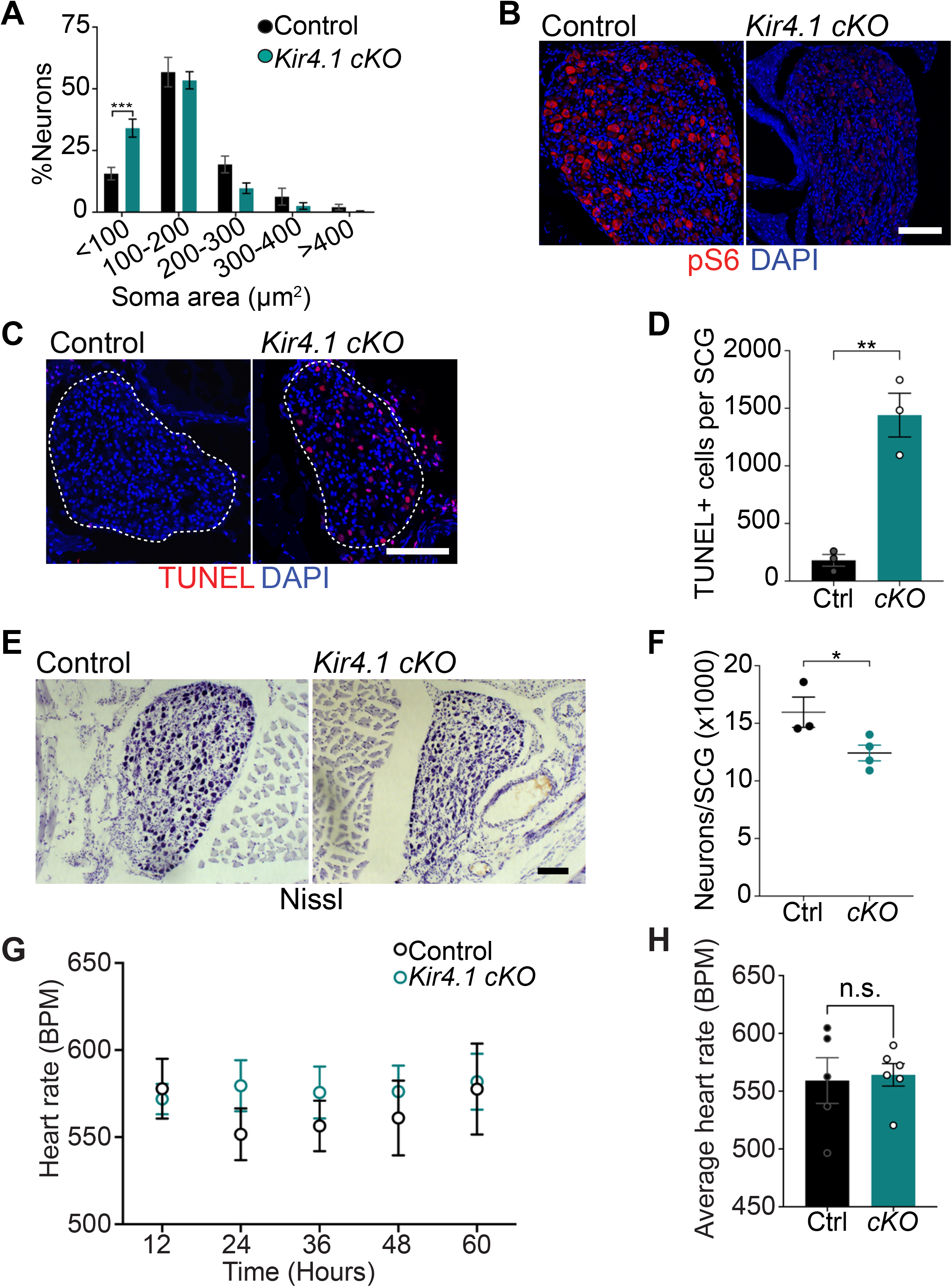
Defects in neuron viability in *Kir4*.*1 cKO* mice. (**A**) *Kir4*.*1 cKO* sympathetic neurons have smaller soma sizes, relative to control neurons. Results are mean ± s.e.m from n=3 control and 5 mutant mice, ***p<0.001, two-way ANOVA with Bonferroni’s correction. (**B**) Decreased mTOR signaling based on pS6 immunostaining in *Kir4*.*1 cKO* sympathetic ganglia. Scale bar:100μm. (**C**) TUNEL labeling shows increased apoptosis in *Kir4*.*1 cKO* SCG (outlined in white dashed line). Nuclei in tissue sections are labeled with DAPI (blue). Scale bar:100μm. (**D**) Quantification of apoptotic cells in SCGs from n=3 mice per genotype. Data presented as mean ± s.e.m. **p<0.01, t-test. (**E-F**) Decreased sympathetic neuron numbers in *Kir4*.*1 cKO* mice based on Nissl-staining and cell counts in SCG tissue sections. Results are mean ± s.e.m from n=3 control and 4 mutant animals. **p<0.01, t-test. (**G**) Heart rate is unaffected by Kir4.1 deletion from satellite glia. Results are mean ± s.e.m from n=5 control and 6 mutant animals, two-way ANOVA with Bonferroni’s correction. (**H**) Average heart rate over days 4-7 post-lead implantation. Results are mean ± s.e.m from n=5 control and 6 mutant animals. n.s., not significant.

We next asked if satellite glia-specific Kir4.1 deletion recapitulates the autonomic defects observed in *BLBP:iDTA* mice. Measurements of basal pupil size, heart rate, and circulating norepinephrine levels indicated that these parameters were not significantly altered in *Kir4*.*1 cKO* mice (**Figures 5G-H, and S5A, B, and D**), despite increased neuronal activity as assessed by c-Fos immunoreactivity (**see Figures 4E-F**). Parasympathetic activity assessed by pupil constriction in response to different light intensities was also normal in *Kir4*.*1 cKO* mice (**Figure S5C**).

Together, these results suggest that Kir4.1 deletion recapitulates the cellular phenotypes observed in *BLBP:iDTA* mice, notably, increased neuron activity, impaired metabolic signaling, and enhanced apoptosis. However, the loss of Kir4.1 is not sufficient to elicit elevated sympathetic tone at the whole-body level as seen with satellite glia depletion.

## Discussion

Despite decades of research on the sympathetic nervous system, satellite glia, the major glial cells in sympathetic ganglia, have remained an enigmatic component of the system. Given their specific contacts with neuronal cell bodies, satellite glia also provide the rare opportunity to study how glia support somatic compartments. Here, we show that satellite glia are essential modulators of sympathetic neuron metabolism, survival, neurotransmitter homeostasis, activity, and autonomic functions in adult mice. Together, our findings provide *in vivo* evidence that neurons and their surrounding glial covers are functional units in the regulation of sympathetic output.

We reveal that a key role for satellite glia is in restraining neuronal activity in mature sympathetic neurons. Depletion of satellite glia in adult mice amplifies neuronal activity leading to increased circulating levels of NE and elevated sympathetic tone. Glia-ablated mice show enhanced pupil size and heart rate, demonstrating the necessity of these cells in the dynamic regulation of autonomic functions in conscious and freely moving animals. Our findings that satellite glia limit neuronal activity are similar to reported functions of other glial cells that encapsulate neuronal cell bodies, in particular, astrocytes (Allen and Lyons, 2018), microglia (Badimon et al., 2020; Cserep et al., 2020), and cortex glia in *Drosophila* (Yadav et al., 2019). Inhibitory effects of satellite glia might enable sympathetic neurons to respond to a wider range of input strengths and/or serve as a neuroprotective mechanism to limit neurotoxicity under conditions of stress or pathology.

How do satellite glia limit sympathetic neuron activity? The enhanced neuronal c-Fos signals in *Kir4*.*1 cKO* and satellite glia-ablated mice, suggest that glial regulation of ion homeostasis, specifically, K^+^ clearance, is a key mechanism that contributes to inhibition of neuronal activity. Satellite glia take up extracellular K^+^, distribute them throughout the glial syncytium via gap junction coupling, and extrude ions in regions of low K^+^ concentration, in a process called “spatial K^+^ buffering” (Kuffler, 1967). Even slight elevations in extracellular K^+^ in the neuronal micro-environment are likely to elicit neuronal depolarization and activation (Cui et al., 2018; Haydon, 2001). However, in contrast to satellite glia-ablated mice, Kir4.1 loss is not sufficient to compromise autonomic function, suggesting that additional glia-dependent mechanisms contribute to influencing neuronal excitability to drive network-level changes. Satellite glia could influence neuronal activity via regulation of ganglionic levels of acetylcholine, the major neurotransmitter released by pre-ganglionic axons. Despite previous observations that satellite glia modulate cholinergic neurotransmission in co-cultured sympathetic neurons (Enes et al., 2020; Feldman-Goriachnik et al., 2018), we did not detect glial transcripts involved in cholinergic signaling using single-cell RNA sequencing of sympathetic ganglia (Mapps et al., 2021). However, we do not exclude glia-mediated effects on synapse formation during development or maintenance (Enes et al., 2020). Another potential mechanism could involve glial regulation of extracellular ATP and/or its break-down products in the neuronal micro-environment (Hanani, 2010). In sympathetic ganglia, ATP is largely released by innervating cholinergic pre-ganglionic axons and facilitates fast excitatory neurotransmission in post-ganglionic sympathetic neurons (Boehm, 1999; Evans et al., 1992; McCaman and McAfee, 1986; Vizi et al., 1997). Extracellular ATP is rapidly metabolized by cell-surface ectonucleotidases, which are expressed in sympathetic satellite glia (Forsman and Elfvin, 1984; Nacimiento et al., 1991; Vizi et al., 1997). Single-cell RNA sequencing data revealed several transcripts involved in ATP sensing, hydrolysis, and removal of break-down products in sympathetic satellite glia, including the purinergic receptors, *P2rx4, P2rx7*, and *P2ry12*, and *Entpd1*, an ectoenzyme which catalyzes ATP to ADP hydrolysis, as well as *Adk* (*adenosine kinase*) (Mapps et al., 2021), which is best-known for mediating astrocytic uptake of extracellular adenosine in the brain (Boison et al., 2010).

We used *BLBPcre-ER* mice in an inducible manner in adult mice, which allowed us to bypass the targeting of cell types during development. BLBP (also called *Fabp7*) is also expressed in neuronal progenitors, radial glia, and astrocytes in the CNS (Ebrahimi et al., 2016; Feng et al., 1994; Matsumata et al., 2012). While we cannot completely exclude an astrocyte-mediated contribution to autonomic defects in *BLBP:iDTA* mice, these mice had normal body weights and no motor deficits, in contrast to mice where adult astrocyte ablation elicited rapid onset of severe limb paralysis and smaller body weights (Schreiner et al., 2015). Of note, our single-cell RNA sequencing analysis of satellite glia (Mapps et al., 2021) compared to published single-cell RNA studies of astrocytes (Batiuk et al., 2020), indicate a ∼45-fold enrichment of *Fabp7* transcript in satellite glia relative to astrocytes. Further, astrocyte-specific deletion of Kir4.1 using *GFAP-cre* transgenic mice results in premature lethality, epileptic seizures, and severe ataxia (Djukic et al., 2007), none of which were observed with Kir4.1 deletion using *BLBPcre-ER* mice. Together, these results suggest that astrocytes are minimally perturbed in *BLBP:iDTA* and *Kir4*.*1 cKO* mice.

In a previous study, acute chemo-genetic activation of satellite glia increased heart rate and cardiac output in *Gfap-hM3Dq* mice (Xie et al., 2017). Sympathetic satellite glia, very likely, exert both excitatory and inhibitory effects on neuronal activity in a context-dependent manner, similar to astrocyte functions in the brain (Allen and Lyons, 2018). Specifically, under conditions of nerve injury or inflammation, activated sensory satellite glia are known to undergo structural and functional changes, resulting in neuronal hyper-excitability (Hanani and Spray, 2020). Limited studies, so far, suggest that in response to nerve damage, sympathetic satellite glia are also capable of reactive changes, specifically, enhanced gap junction-mediated coupling and increased ATP sensitivity (Feldman-Goriachnik and Hanani, 2019), which in turn, may augment neuronal activity.

An intriguing finding was that satellite glia depletion or glial deletion of *Kir4*.*1* resulted in the loss of ∼25% of adult sympathetic neurons. The exquisite dependence of sympathetic neurons on the target-derived survival factor, Nerve Growth Factor (NGF), during development is well-documented (Glebova and Ginty, 2005). However, loss of NGF signaling does not compromise the survival of adult neurons (Angeletti et al., 1971; Tsui-Pierchala and Ginty, 1999), and to date, the trophic mechanisms underlying adult sympathetic neuron survival *in vivo* remain undefined. The similar phenotypes of soma atrophy, impaired mTOR signaling, and enhanced apoptosis in adult *BLBP:iDTA* and *Kir4*.*1 cKO* sympathetic neurons suggest that satellite glia provide metabolic and trophic support to mature neurons, primarily, via regulation of K^+^ homeostasis. Of note, astrocyte-specific *Kir4*.*1* deletion also results in defects in mTOR signaling and decreased soma size, but not cell death, in a population of spinal cord motor neurons (Kelley et al., 2018). Whether observed apoptosis of mature sympathetic neurons in *BLBP:iDTA* or *Kir4*.*1 cKO* mice is linked to hyper-excitability of a vulnerable neuronal sub-population and/or reflects the requirement for glial-derived trophic signals remain to be defined. Further, despite enhanced sympathetic neuron activity and circulating NE levels in satellite glia-ablated mice, we observed a pronounced down-regulation of NE biosynthetic machinery, specifically in neuronal cell bodies. This may reflect a compensatory response to elevated sympathetic activity upon glia loss or the need for glia-derived factors in maintenance of noradrenergic enzymes. It is also notable that satellite glia depletion resulted in soma-specific effects in sympathetic neurons, i.e., TH down-regulation and soma size decrease, without perturbing axonal TH levels and innervation. Together, these findings highlight the compartment-specific functions of satellite glia-neuron interactions.

Heightened activity of the sympathetic nervous system is a characteristic feature of several pathological conditions, specifically, chronic heart failure (Malpas, 2010), arrhythmias (Hasan, 2013; Schwartz, 2014), hypertension (Goldstein et al., 2002; Schlaich et al., 2004), sleep apnea, obesity, and insulin resistance (Mahfoud et al., 2011; Mancia et al., 2007; Schlaich et al., 2004). While sympathetic neurons, so far, have been at the center stage in considering pathological mechanisms and treatments, our study, together with other recent work (Xie et al., 2017), highlight the therapeutic potential of targeting the neuron-satellite glia unit in autonomic-related diseases.

## STAR METHODS

- KEY RESOURCES TABLE
- CONTACT FOR REAGENT AND RESOURCE SHARING
- EXPERIMENTAL MODEL AND SUBJECT DETAILS
  - Animals
- METHOD DETAILS
  - Tamoxifen injections
  - qRT-PCR
  - smFISH
  - Immunohistochemical analyses
  - IDISCO whole-mount immunostaining
  - Soma size
  - Neuronal cell counts
  - TUNEL
  - EdU labeling
  - Norepinephrine ELISA
  - Pupil analyses
  - Electrocardiograms

QUANTIFICATION AND STATISTICAL ANALYSES

## Acknowledgements

We thank all members of the Kuruvilla, Zhao, and Hattar labs for helpful comments on the project. This work was supported by NIH R01 awards, NS073751 and NS107342, to R.K., DC016065 and EY027202 to H.Z, NIMH intramural research funds (ZIAMH002964) to S.H., a NSF GRFP (DGE-1746891) award to A.M., and NIH Training grant T32GM007231 to the JHU CMDB graduate program for A.M and E.B.

## Author Contributions

A.M. was primarily responsible for the study design, investigations, data analyses, and contributed to writing and editing the manuscript. E.B., C.B., J.L., W.K., M.L., and E.T., assisted with experiments. R.K, H.Z, and S.H contributed to writing/editing the manuscript and funding acquisition.

## STAR METHODS

### KEY RESOURCES TABLE

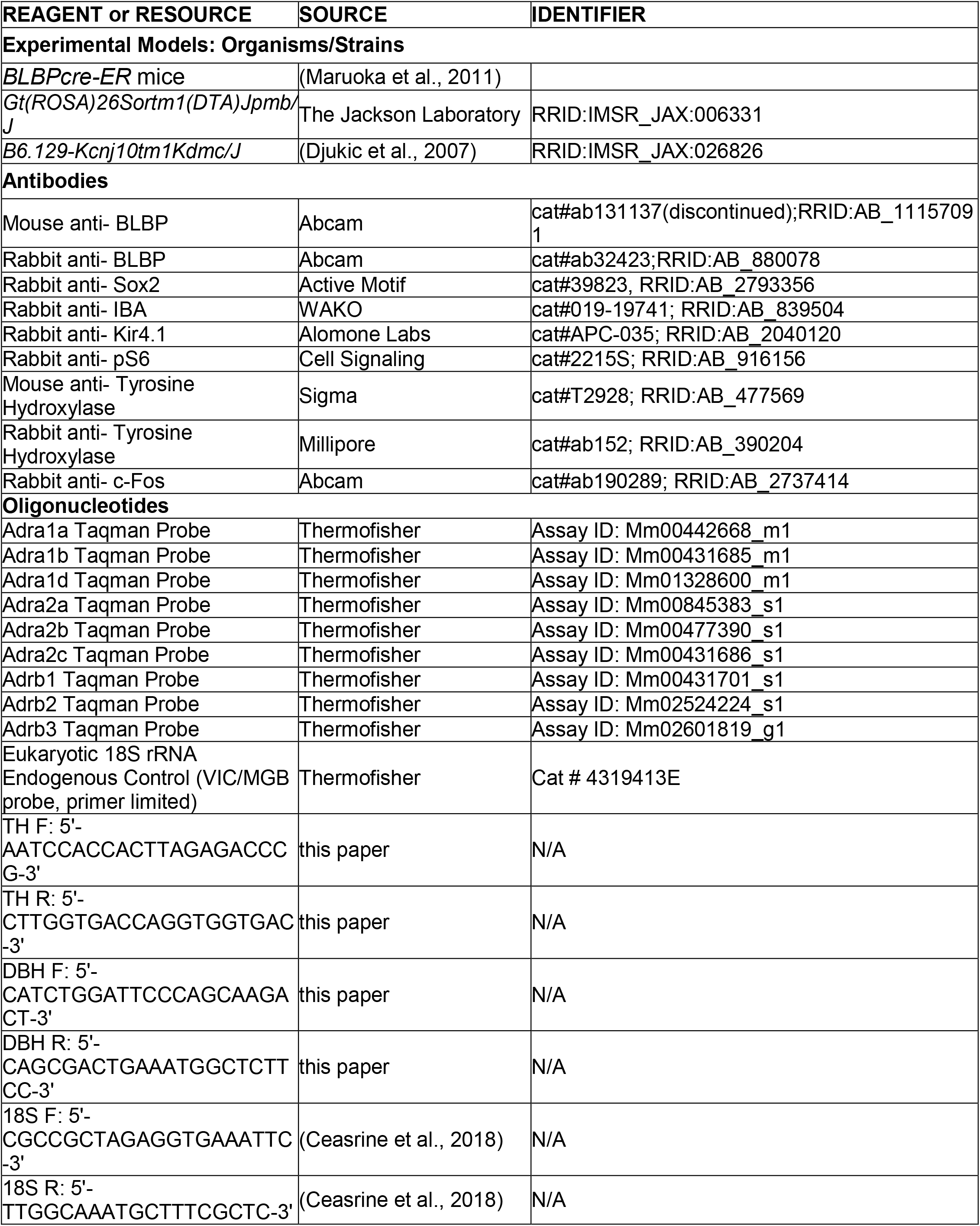
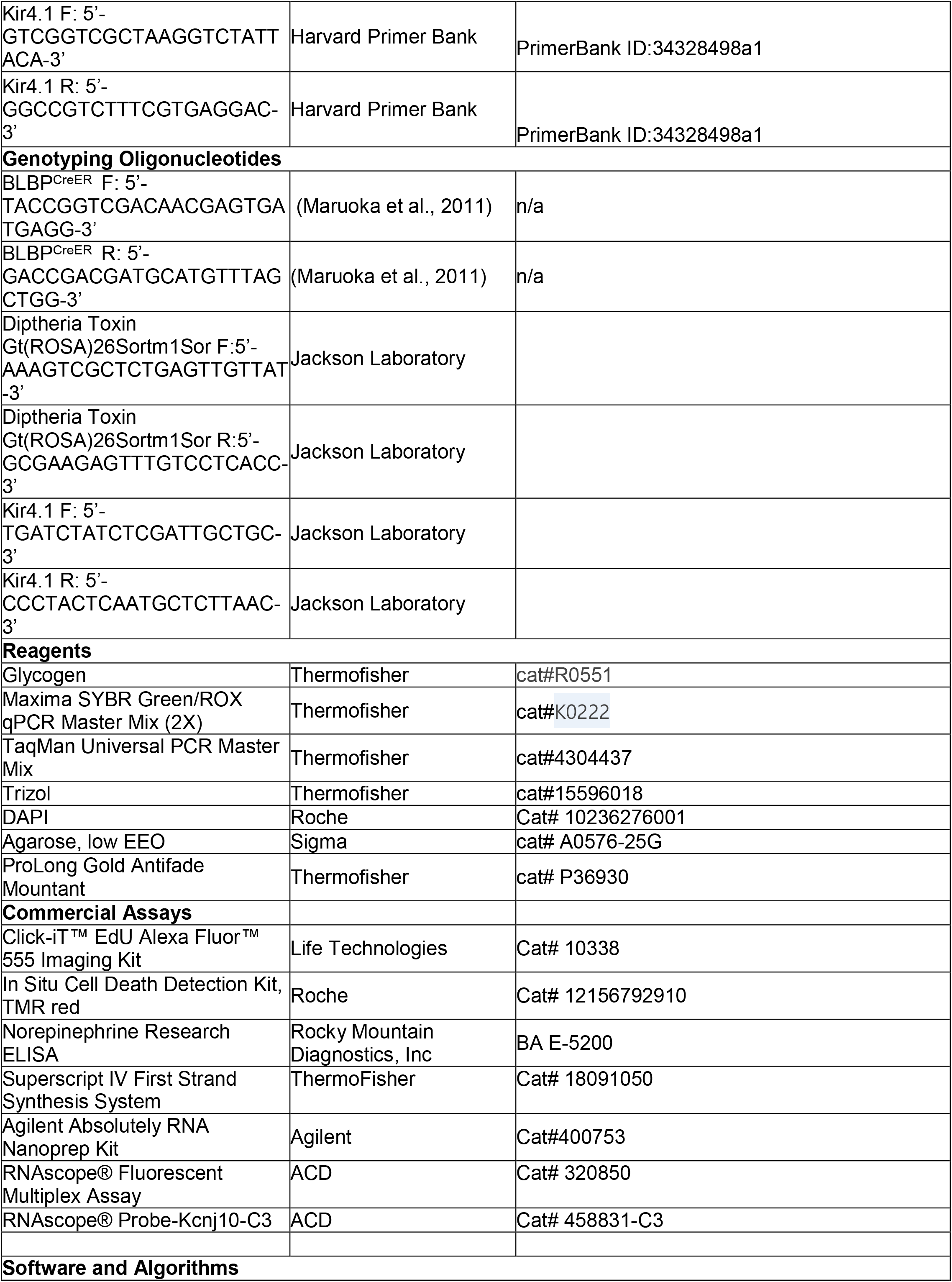
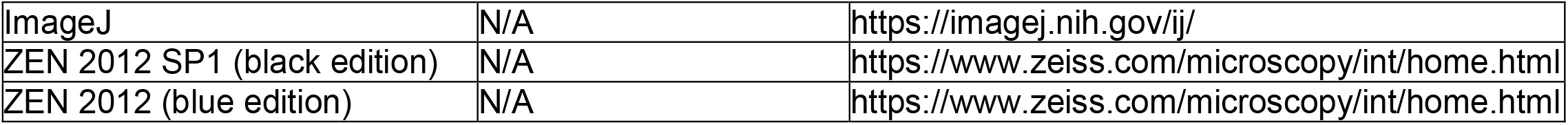

### CONTACT FOR REAGENT AND RESOURCE SHARING

Further information and requests for resources and reagents should be directed to and will be fulfilled by the Lead Contact, Rejji Kuruvilla.

#### Data and Code Availability

This study did not generate/analyze datasets/code.

### EXPERIMENTAL MODEL AND SUBJECT DETAILS

#### Animal care and housing

All procedures relating to animal care and treatment conformed to The Johns Hopkins University Animal Care and Use Committee (ACUC) and NIH guidelines. Animals were group housed in a standard 12:12 light-dark cycle, except for the pupil analyses. Adult mice between 1-1.5 months of age, and of both sexes, were used for analyses. The following mouse lines were used in this study: *BLBPcre-ER* mice (Maruoka et al., 2011) were generously provided by Dr. Toshihiko Hosoya (RIKEN Brain Science Institute), *Kir4*.*1*^*fl/fl*^ mice (Djukic et al., 2007) (Jackson Laboratory, stock no: 026826) by Dr. Dwight Bergles (Johns Hopkins School of Medicine). *ROSA26-eGFP-DTA* mice were obtained from the Jackson Laboratory (stock no: 006331).

### MATERIALS & METHODS

#### Tamoxifen injections

At postnatal day 30, *BLBPcre-ER*;*ROSA26-eGFP-DTA* or *BLBPcre-ER;Kir4*.*1*^*fl/fl*^ mice were injected subcutaneously with either vehicle corn oil (Sigma) or tamoxifen (180 mg/kg body weight) dissolved in corn oil for 5 consecutive days. All analyses were performed at 5 or 14 days after the last injection.

#### qRT-PCR

RNA was isolated from dissected SCGs or cardiac tissue using Absolutely RNA Nanoprep Kit (Agilent) or Trizol-chloroform extraction. cDNA was prepared using Superscript IV First Strand Synthesis System. Real-time qPCR analysis was performed using Maxima SYBER Green/Rox Q-PCR Master Mix (Thermo-Fisher) and gene specific primers for SCG tissues or Taq-Man probes for adrenergic receptors in heart tissue, in a 7300 Real time PCR System (Applied Biosystems) Each sample was analyzed in triplicate reactions. Fold change was calculated using the 2^(ΔΔ*Ct*)^ method, normalizing to 18S transcript.

#### smFISH

SCGs were dissected from P48 mice, cryo-protected in 30% sucrose in PBS for 1 hr and embedded in OCT and frozen at −80°C. Ganglia were cryo-sectioned at 14 µm and kept at −80°C until smFISH was performed. Target mRNA was probed using RNAscope® Multiplex Fluorescent Reagent Kit v2 Assay. Tissues were incubated in fresh 4% paraformaldehyde for 5 min, washed twice in 1xPBS, and dehydrated with increasing concentrations of ethanol. Subsequently, tissues were treated with hydrogen peroxide for 10 min and protease treatment for 15 min. RNAscope assays were performed following the manufacturer’s instructions. Tissues were mounted with Prolong Anti-fade Mounting Media, and imaged using a Zeiss LSM 800 confocal microscope.

#### Immunohistochemical analyses

Superior cervical ganglia (SCG) were harvested and incubated in 4% paraformaldehyde for 4 hr at room temperature, cryo-protected in 30% sucrose in 1xPBS for 3 days, embedded in OCT (Sakura Finetek) and stored at −80°C. Ganglia were cryo-sectioned at 12-30μm sections for immunohistochemistry. For paraffin embedding, SCGs were fixed in Bouin’s solution for 1 hr at room temperature, left in 70% ethanol overnight at room temperature, followed by consecutive washes in 70%, 80%, 95%, and 100% ethanol, and xylene. Tissues were embedded in paraffin wax and sectioned at 6µm using a microtome, and rehydrated using a series of xylene and ethanol washes at room temperature. Tissue sections were permeabilized in 0.1% triton x-100 in 1x PBS at room temperature 3X for 5 min each, followed by incubation in 10mM sodium citrate buffer (pH 6) at 95°C for 10 min. Tissues were blocked in 5% goat serum/3% bovine serum albumin in 0.1% triton x-100 in 1xPBS (blocking solution) for 1 hr at room temperature. Primary antibodies used were mouse anti-BLBP (1:500), rabbit anti-BLBP (1:200), mouse anti-TH (1:300), rabbit anti-TH (1:300), rabbit anti-IBA1 (1:200), rabbit anti-pS6 (1:200), rabbit anti-c-Fos (1:1000), or rabbit anti-Kir4.1 (1:100) with incubations performed overnight at 4°C. Slides were washed in blocking solution and then incubated in Alexa-546 or-647 conjugated anti-rabbit or anti-mouse secondary antibodies (1:200 dilution) and DAPI (1:1000) in blocking buffer. Tissues were then mounted in Aqueous Mounting Medium and imaged at 1µm optical slices using a Zeiss LSM 700 or LSM 800 confocal microscope. Maximum intensity projections were generated with Image J (Fiji).

For Sox-2 immunostaining, SCGs were embedded in 3% agarose (Sigma) and sectioned at 100µm thick sections using a vibratome. Tissues were permeabilized/blocked in 10% goat serum/3% tween in 1x PBS for 2 hr at room temperature, incubated with rabbit anti-Sox2 (1:500) and mouse anti-TH (1:300) antibodies in blocking solution for 2 days at 4°C, followed by Alexa-546 or-647 conjugated anti-rabbit or anti-mouse secondary antibodies (1:400 dilution) and DAPI (1:1000). Tissues were then processed for confocal imaging as stated above.

#### iDISCO and whole mount immunostaining

iDISCO-based tissue clearing for whole mount immunostaining of organs from P48 mice was performed as previously described (Renier et al., 2014). Briefly, hearts were fixed in 4%PFA/PBS, then dehydrated by methanol series (20-80%) and incubated overnight in 66% dichloromethane (DCM)/33% methanol. Samples were then bleached with 5% H_2_O_2_ in methanol at 4°C overnight, then re-hydrated and permeabilized first with 0.2%TritonX-100 followed by overnight permeabilization with 0.16% TritonX-100/20%DMSO/0.3M glycine in PBS. Samples were incubated in blocking solution (0.17% TritonX-100/10% DMSO/6% Normal Goat Serum in PBS) for 8 hr, and then incubated with rabbit-anti-TH (1:400) in 0.2% Tween-20/0.001% heparin/5% DMSO/3% Normal Goat Serum in PBS at 37°C for 96 hr. Samples were then washed with 0.2% Tween-20/0.001% heparin in PBS and incubated with anti-rabbit Alexa-546 secondary antibody (1:400) in 0.2% Tween-20/0.001% heparin/3% Normal Goat Serum in PBS. After 96 hr, organs were extensively washed with 0.2% Tween-20/0.001% heparin in PBS and dehydrated in methanol. Samples were cleared by successive washes in 66% DCM/ 33% methanol, 100% DCM and 100% Dibenzyl Ether. Organs were imaged on a lightsheet microscope (LaVision BioTec Ultra Microscope II). Imaris was used for 3D manipulations. Total axon lengths and number of branch points were quantified using Imaris Filament Tracer and normalized to total organ volume.

#### Soma Size

SCG tissue sections were labeled with hematoxylin and eosin. Cells with characteristic neuronal morphology and visible nucleoli were identified, soma were traced and areas (µm^2^) quantified using ImageJ (FIJI).

#### Neuronal Cell Counts

Neuronal counts were performed as previously described (Scott-Solomon and Kuruvilla, 2020). In brief, torsos of P39-48 mice were fixed in 4%PFA/PBS overnight and cryoprotected in 30% sucrose/PBS for 48 hr. Torsos were then mounted in OCT and serially sectioned (12 μm). Next, every 5th section was stained with solution containing 0.5% cresyl violet (Nissl). Cells in both SCGs with characteristic neuronal morphology and visible nucleoli were counted using ImageJ.

#### TUNEL

Apoptotic cells were identified in every 5th section of ganglia. Tissues were first incubated with primary and secondary antibodies as described above, followed by detection of cell death using TUNEL staining (Roche) according to the manufacturer’s protocol. Cells that were double positive for DAPI and TUNEL were counted as dying cells.

#### EdU labeling

Tamoxifen- or corn oil-injected adult mice were injected intraperitoneally with EdU (Invitrogen, 100 µg/ml in 3:1 PBS/DMSO) for 5 consecutive days before harvesting at 5 days post tamoxifen injections. SCG tissue sections (12 μm) were processed for 30 min at room temperature in EdU reaction cocktail (ThermoFisher EdU kit C10337; Click-iT buffer, Buffer additive, CuSO4 solution, and Alexa-Fluor 488). Sections were then washed in PBS + 0.1% TritonX-100 and mounted with Fluoromount +DAPI. Images were collected using a Zeiss LSM 700 confocal microscope. The total number of cells that incorporated EdU in each section were counted and summed for an entire SCG.

#### Norepinephrine ELISA

Blood samples drawn from anesthetized mice were centrifuged for 15 min, 3000 rpm in 0.5M EDTA-coated tubes at 4°C. Norepinephrine levels in plasma (300μl) was assessed using a ELISA kit (ABNOVA) according to the manufacturer’s protocol.

#### Pupil analyses

Pupil size measurements were performed as reported previously (Keenan et al., 2016). Briefly, all mice were dark-adapted and housed in single cages for 2 days and analyzed in the evenings. For all experiments, mice were un-anesthetized and restrained by hand. To mitigate stress, which can affect pupil size, researchers handled mice for several days prior to the measurements. Videos of the eye were recorded for 5 to 10 sec in the dark using a Sony Handycam (DCR-HC96) mounted on a tripod at a fixed distance from the mouse. Manual focus was maintained on the camera to ensure that only one focal plane existed for each mouse. Pupil size was recorded under dim red light and the endogenous infrared light source of the camera to capture the basal pupil size.

To examine parasympathetic activity, mice were dark-adapted for 2 days. Un-anesthetized mice were restrained by hand. Pupil size was recorded first for 5 to 10 sec in the dark followed by a 30 sec exposure to a light step stimulus. The light stimulus was provided by 10W or 14W LED Bulbs (Sunlite A19/ or Sunlite 80599-SU LED A19 Super Bright Light Bulb, Daylight). Light intensity was measured using a light meter (EXTECH Foot Candle/Lux Light Meter, 401025) at the surface on which the mouse was held. Light intensity was adjusted by a combination of altering the distance of the light bulb from the mouse and/or applying neutral density filters (Roscolux). The light meter is unreliable at detecting light intensities below 1 lux, so one neutral density filter cutting the light intensity by 12.5% was applied to the bulb to estimate 1-log unit decreases in illumination below 1 lux. Light intensities above 500 lux required the use of multiple light bulbs.

#### Electrocardiograms

Electrocardiogram (ECG) recordings were performed on adult mice as previously described (Stahlberg et al., 2019). Briefly, adult mice (P39-48) were anesthetized with 4% isoflurane, intubated, and placed on ventilator support (settings 1.2ml/g/min at 80 breaths/minute). The animal’s dorsum was shaved, scrubbed with betadine and alcohol, and draped with a sterile barrier with the surgery site exposed. A small (0.5cm midline incision was performed and ECG leads were implanted subcutaneously and sutured over the trapezius muscle on both sides. Body temperature was maintained at 37°C. Immediately following implantation, the wound was closed with a 3-0 silk suture. Anesthesia was turned off and the animal was monitored for spontaneous breathing and was given a subcutaneous buprenorphine injection (0.01–0.05mg/kg buprenorphine, IM) to alleviate pain. ECGs were subsequently recorded continuously in conscious animals for approximately 7 days for each mouse using the Powerlab data acquisition device and LabChart 8 software (AD instruments). Mice were kept at a stable temperature with regular 12-hour light/dark cycle. To exclude the effects of pain and anesthesia, continuous ECG recordings between day 4-7 post lead implantation were only included in the analysis of mean heart rates. Heart rate variability was analyzed using LabChart 8 and at 10-12-hour intervals as previously described (Thireau et al., 2008).

#### Quantification and statistical analyses

Sample sizes in this study were calculated using power analyses using R Studio statistical software. For practical reasons, analyses of innervation, cell death assays, proliferation assays, soma size measurements, and norepinephrine ELISA’s were performed in a semi-blinded manner. The experimenter was aware of the genotypes but performed each immunostaining and measurements without knowing the genotypes. All physiological experiments (pupil dilation and heart rate variability) were performed in a blinded manner; the experimenter was only aware of the ear tag numbers. All t-tests were performed assuming Gaussian distribution, two-tailed, unpaired and a confidence interval of 95%. One-way or two-way ANOVA analyses with Bonferroni’s correction were performed when more than two groups were compared. All error bars represent the standard error of the mean (s.e.m.).

**Supplemental Figure S1.**
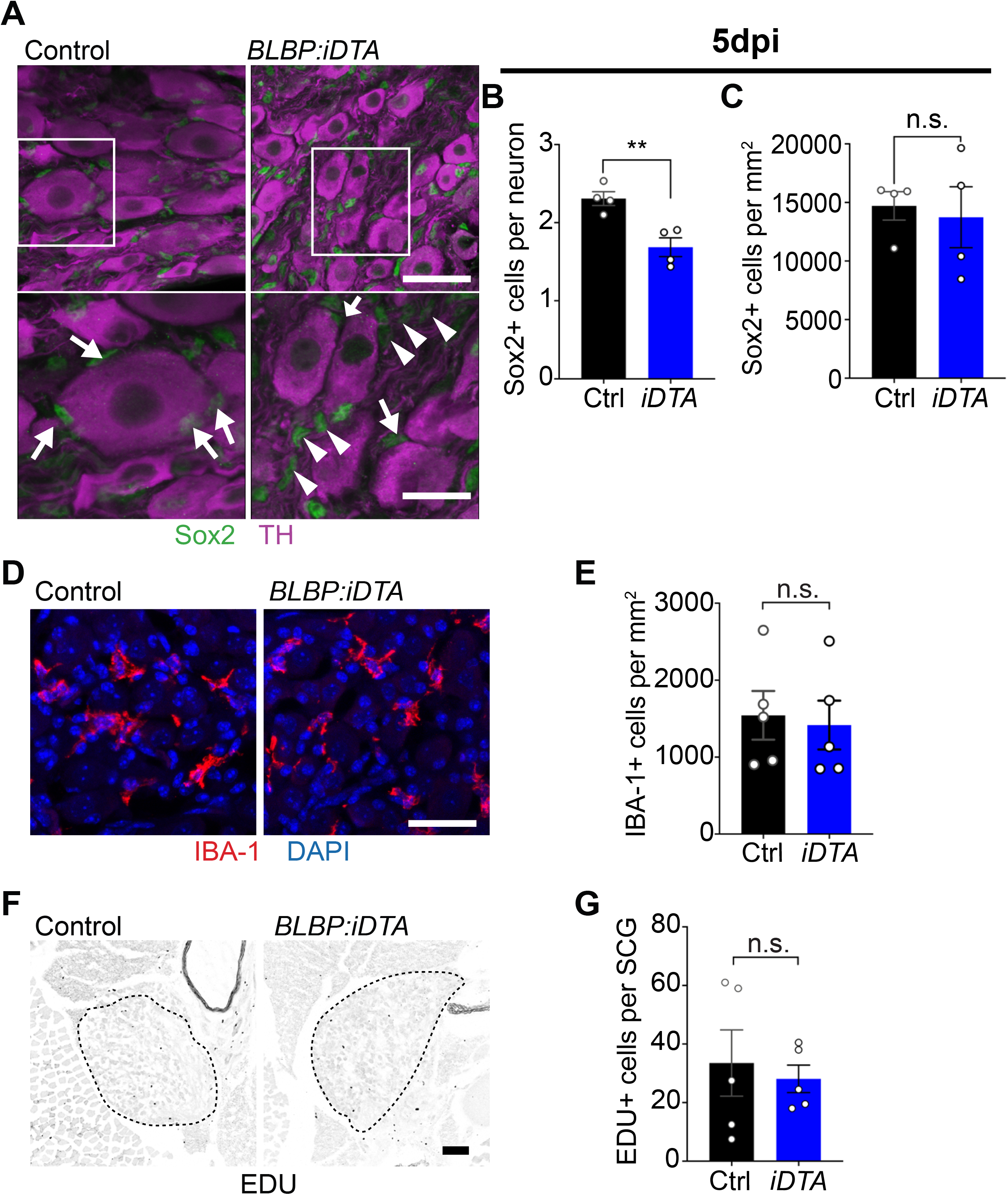
Satellite glia depletion does not induce inflammation or proliferative changes in sympathetic ganglia. **(A-C)** Decreased association of Sox2-positive satellite glia with neuron cell bodies in *BLBP:iDTA* sympathetic ganglia 5 days after last tamoxifen injection (5 dpi). Arrows indicate Sox2-labeled nuclei of satellite glia associated with sympathetic neuron cell bodies. Arrowheads indicate Sox2-labeled satellite glia that are not associated with neurons. Glial cell numbers are unaffected at this time. Scale bar:100μm for upper panels and 30 μm in insets. Data are mean ± s.e.m from n=4 animals per genotype, **p<0.01, n.s., not significant, t-test. (**D-E**) Satellite glia depletion does not induce an increase in IBA-1-expressing macrophages in sympathetic ganglia. Scale bar:30 µm. Data are mean ± s.e.m. from n=5 animals per genotype, n.s., not significant, t-test. (**F-G**) Proliferation in SCGs (outlined by dashed line) was not altered by satellite glia depletion as assessed by EDU labeling. Scale bar:100 µm. Data are mean ± s.e.m. from n=5 animals per genotype, n.s., not significant, t-test.

**Supplemental Figure S2.**
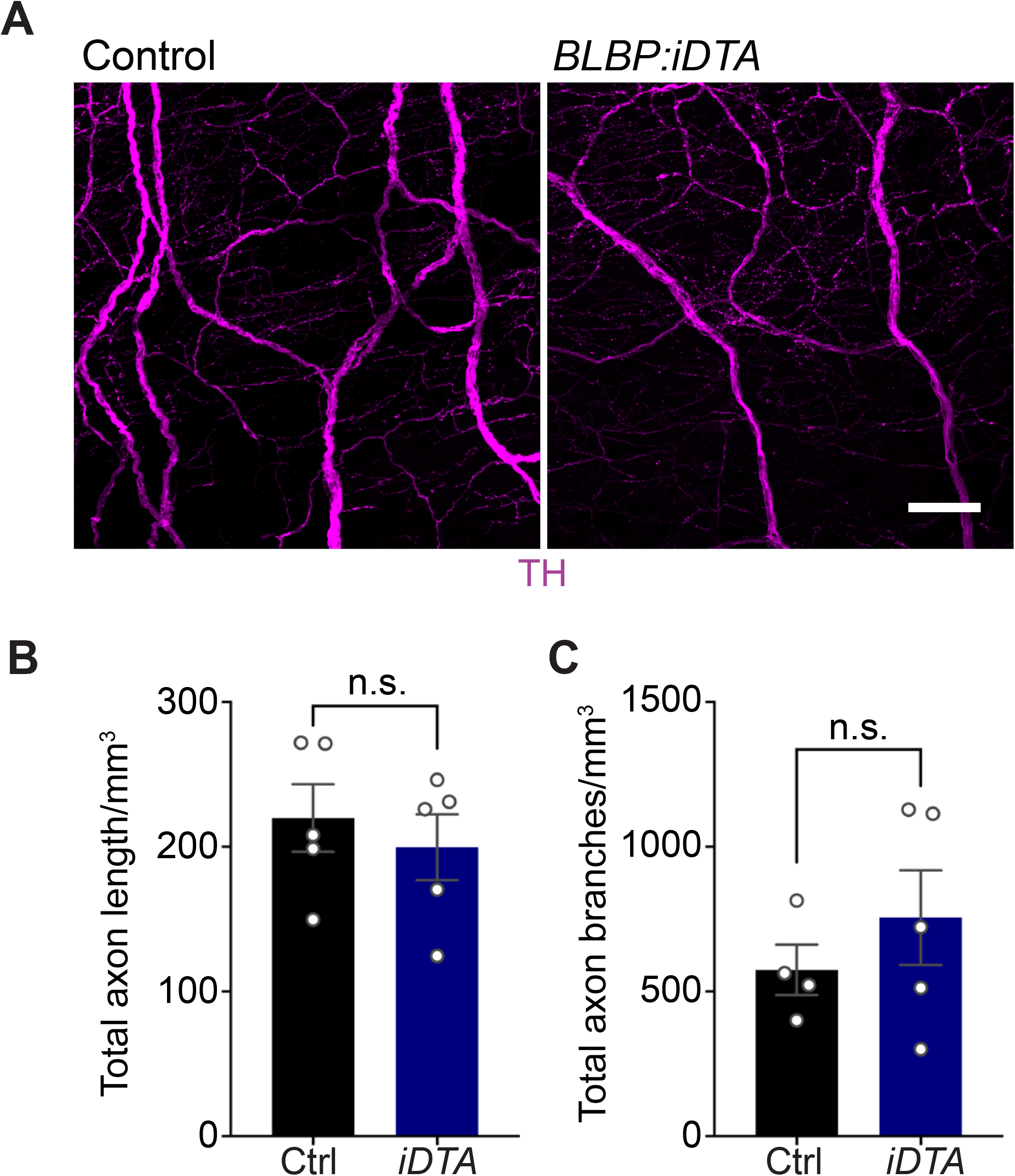
Sympathetic axon innervation of targets is unaffected in satellite glia depleted mice. **(A)** Sympathetic axon innervation of the heart is unaffected in adult *BLBP:iDTA* mice based on iDISCO-based tissue clearing and wholemount TH immunostaining. Scale bar:100µm. (**B-C**) Quantification of axon length and branches in the heart in *BLBP:iDTA* and control mice. Data are mean ± s.e.m. from n=5 animals per genotype, n.s., not significant, t-test.

**Supplemental Figure S3.**
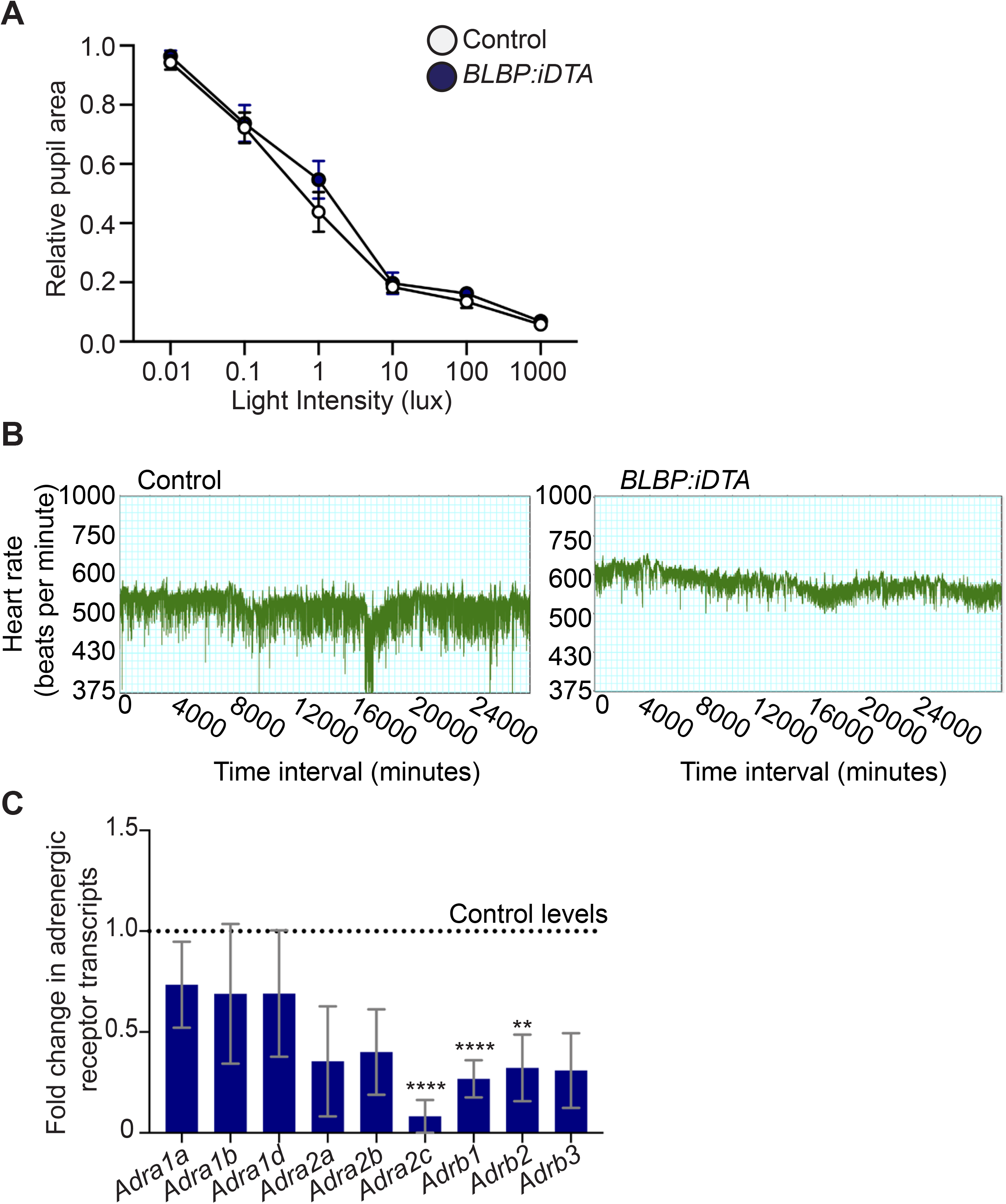
Autonomic function analyses in *BLBP:iDTA* mice. **(A)** Parasympathetic activity is normal in satellite glia-depleted mice as assessed by measuring pupil constriction in response to increasing light intensities. Data are mean ± s.e.m. from n=5 control and 6 mutant animals, two-way ANOVA with Bonferroni’s correction. (**B**) Representative heart rate recordings over time show decreased heart rate variability in *BLBP:iDTA* mice compared to controls. **(C)** Down-regulated adrenergic receptor expression in cardiac tissue as assessed by qPCR analysis. Data are mean ± s.e.m. from n=6 control and 5 mutant animals, **p<0.01, ***p<0.001, ****p<0.0001, t-test with Bonferroni-Dunn’s correction.

**Supplemental Figure S4.**
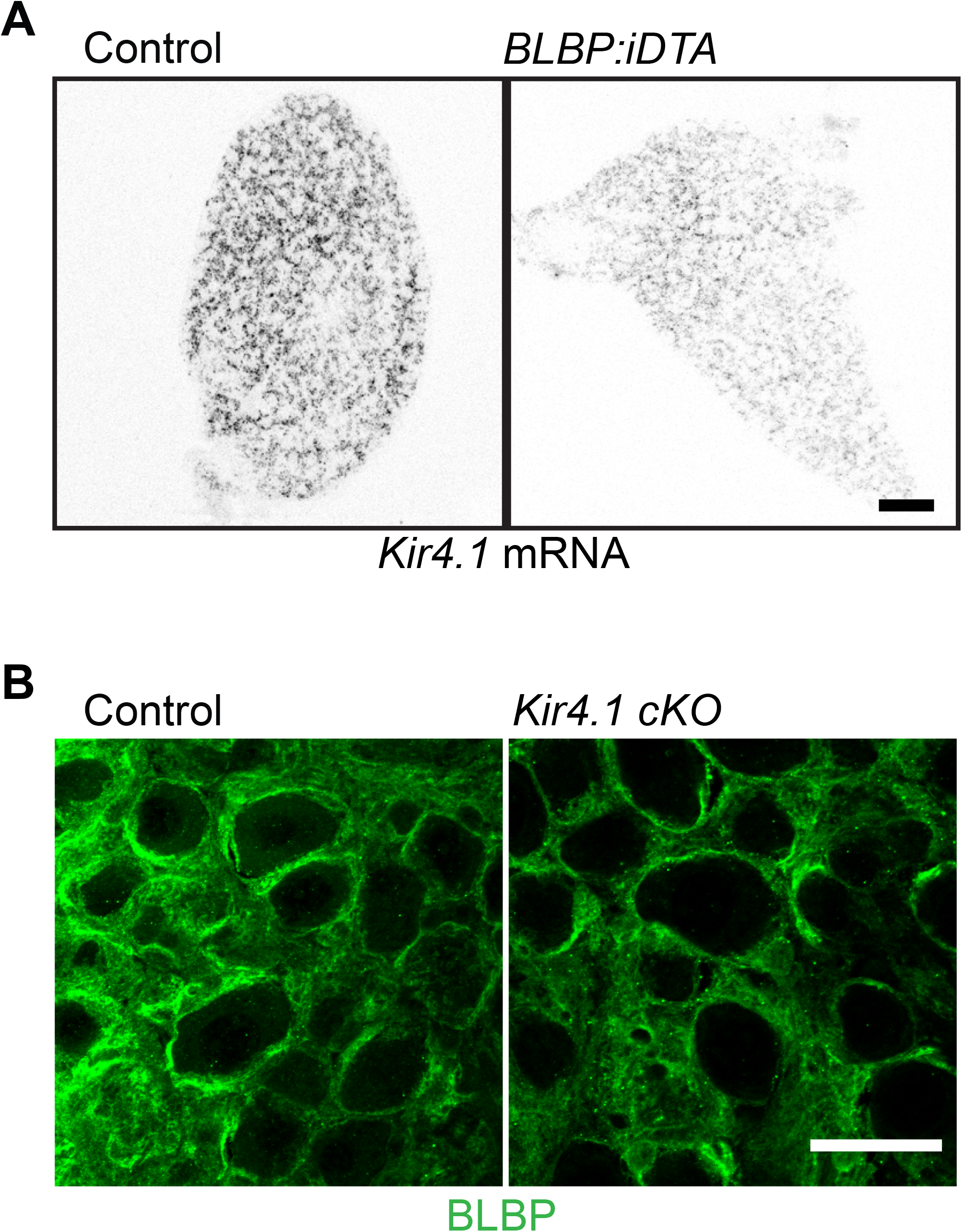
*Kir4*.*1* and BLBP expression in satellite glia-deficient mice. **(A)** smFISH shows decreased *Kir4*.*1* mRNA in BLBP:iDTA sympathetic ganglia. Scale bar: 100µm. **(B)** BLBP immunostaining shows that satellite glial morphologies and BLBP expression are not altered in *Kir4*.*1 cKO* sympathetic ganglia. Scale bar: 30µm.

**Supplemental Figure S5.**
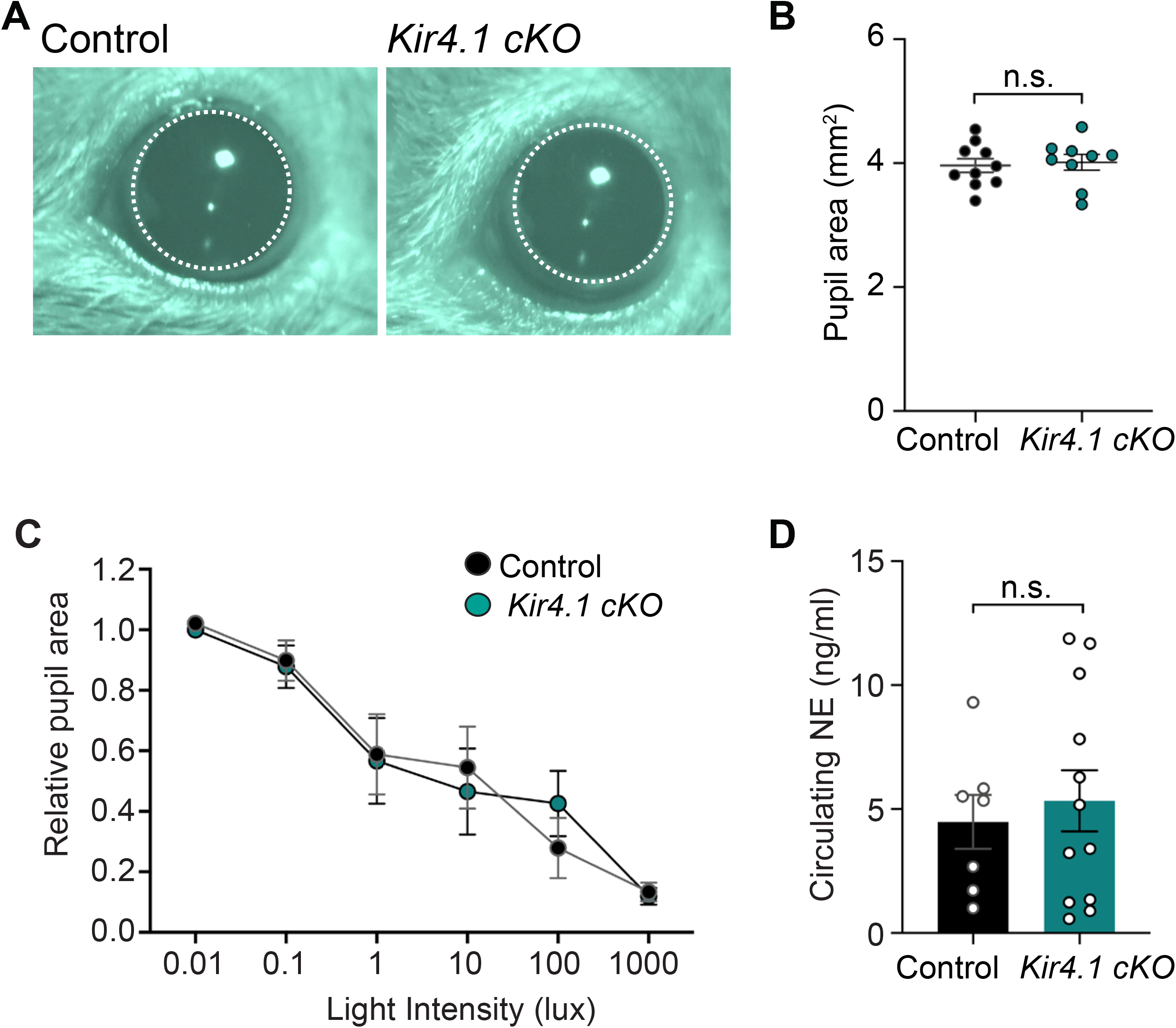
Autonomic function analyses in *Kir4*.*1 cKO* mice. **(A-B)** Basal pupil areas are unaffected by satellite glia-specific Kir4.1 deletion. (**C**) Pupil constriction in response to light, indicative of parasympathetic activity, is normal in *Kir4*.*1 cKO* mice. Data are mean ± s.e.m. from n=6 animals per genotype, two-way ANOVA with Bonferroni’s correction. (**D**) Circulating norepinephrine (NE) levels are unaltered with Kir4.1 deletion from satellite glia. Data are mean ± s.e.m. from n=7 control and 12 mutant animals, n.s. not significant, t-test.

